# Correlated evolution between heat tolerance and thermal performance curves in *Drosophila subobscura*

**DOI:** 10.1101/864793

**Authors:** Andrés Mesas, Angélica Jaramillo, Luis E. Castañeda

**Affiliations:** Laboratorio de Genómica y Biodiversidad, Departamento de Ciencias Básicas, Facultad de Ciencias, Universidad del Bío-Bío, Chillán, Chile; Programa de Genética Humana, Instituto de Ciencias Biomédicas, Facultad de Medicina, Universidad de Chile, PO 8380453, Santiago, Chile

**Keywords:** artificial selection, climate change, experimental evolution, heat tolerance evolution, locomotor performance, thermal reaction norm, thermal stress

## Abstract

Global warming imposes important challenges for ectotherm organisms, which can avoid the negative effects of thermal stress via evolutionary adaptation of their upper thermal limits (CT_max_). In this sense, the estimation of CT_max_ and its evolutionary capacity is crucial to determine the vulnerability of natural populations to climate change. However, these estimates depend on the thermal stress intensity and it is not completely clear whether this thermal stress intensity can impact the evolutionary response of CT_max_ and thermal reaction norms (i.e. thermal performance curve, TPC). Here we performed an evolutionary experiment by selecting high heat tolerance using acute and chronic thermal stress in *Drosophila subobscura*. After artificial selection, we found that knockdown temperatures (a CT_max_ proxy) evolved in selected lines compared to control lines, whereas the realized heritability and evolutionary rate change of heat tolerance did not differ between acute-selected and chronic-selected lines. From TPC analysis, we found acute-selected lines evolved a higher optimal performance temperature (T_opt_) compared to acute-control lines, whereas this TPC parameter was not different between chronic-selected and chronic-control lines. The evolutionary response of T_opt_ caused a displacement of entire TPC to high temperatures suggesting a shared genetic architecture between heat tolerance and high-temperature performance, which only arose in the acute-selected lines. In conclusion, thermal stress intensity has important effects on the evolution of thermal physiology in ectotherms, indicating that different thermal scenarios conduce to similar evolutionary responses of heat tolerance but do not for thermal performance. Therefore, thermal stress intensity could have important consequences on the estimations of the vulnerability of ectotherms to global warming.

## Introduction

Environmental temperatures influence the organismic functions and fitness of ectotherm animals (Huey & Stevenson, 1979; Huey & Kingsolver, 1993; Angilletta et al., 2009), causing that distribution and abundance of these organisms to be mainly limited by their thermal limits (Parmesan & Yohe, 2003; Beugrand et al., 2002; Lima et al., 2007; Pannetta et al., 2018). Thus, the estimation of the evolutionary response of upper thermal limits (CT_max_) it is crucial to predict and understand the capability of ectotherms to respond to increasing thermal challenges (Gunderson & Stillman, 2015). In this context, several studies have revealed a limited evolutionary potential of CT_max_ in ectotherms, suggesting that a high vulnerability of ectotherms to global warming (Sunday et al., 2011; Kellerman et al., 2012; Kelly et al., 2012; Araujo et al., 2013; Hoffmann et al., 2013). However, some experimental studies have reported a large evolutionary response for CT_max_ only after a few generations under selection for increasing thermotolerance (Bubliy & Loeschcke, 2005; Folk et al., 2006; Geerts et al, 2015; Hangartner & Hoffmann, 2016). Consequently, it is mandatory to elucidate the factors that could explain this contrasting evidence about the heat tolerance evolution.

Commonly, CT_max_ is estimated using dynamic assays, in which individuals are exposed to non-stressful temperatures and then temperature is increased at a specific rate until the organisms collapse (Cowles y Bogert, 1944; Hutchinson, 1961; Lutterschmidt y Hutchison, 1997). However, several lines of evidence suggest that heat tolerance estimates depend on the intensity of thermal stress employed during dynamic assays, showing that organisms exposed to a chronic thermal stress (e.g. slow ramping assays) exhibit a lower CT_max_ than those individuals exposed to an acute thermal stress (e.g. fast ramping assays) (Terblanche et al., 2007; Chown et al., 2009; Peck et al., 2009; Mitchell & Hoffmann, 2010; Ribeiro et al., 2012). This methodological impact on heat tolerance estimates has been explained as the consequences of the physiological mechanisms related to heat stress such as resource depletion, water loss and heat-induced cellular damage, which can be more important during longer thermal assays (Rezende et al. 2011; Kingsolver & Umbanhowar, 2018). Interestingly, the intensity of thermal stress also has effects on the heritability estimates of CT_max_: the longer the thermal assays, the lower the heritability (Mitchell & Hoffmann, 2010; Blackburn et al., 2014; Castañeda et al. 2019). Therefore, these resistance-associated mechanisms (e.g. resource depletion, water loss, cellular damage) are expected to increase the environmental variance of CT_max_ assayed under chronic thermal stress, leading to a reduced heritability estimates in comparison to parameters estimated under acute thermal stress (Chown et al., 2009; Rezende et al. 2011). However, to date the impact of the thermal stress intensity on the evolutionary response of heat tolerance methodology has been only explored through computer simulations by mimicking artificial selection experiments for increasing heat tolerance using different ramping rates in *Drosophila melanogaster* (Santos et al., 2012). In this simulation, Santos et al. (2012) found that not only a reduced increase of heat tolerance in slow-ramping selected lines compared to fast-ramping selected lines was evident, but also that there was a correlated physiological response (e.g. metabolic rate) in slow-ramping selected lines. Then, these simulations suggest that the evolutionary response of heat tolerance and correlated responses depend on the thermal stress intensity, which could be the result of the how much precise is the estimation of CT_max_ (Castañeda et al. 2019). However, there is not empirical evidence that supports this hypothesis to date and it is the main goal of the present work.

The correlated responses to the evolution of CT_max_ may be the result of the genetic architecture underlying heat tolerance (Angilletta, 2009; Hangartner et al., 2016; Rolandi et al., 2018), but the impact of the thermal stress intensity on the correlated evolutionary response of other phenotypic traits related to CT_max_ is unknown. In their seminal work, Huey & Kingsolver (1993) proposed different evolutionary consequences of the thermal performance curve (TPC) as response to selection for increased heat tolerance. The TPC is a reaction norm that describes the complex relationship between the organismic performance (e.g. locomotion, growth, development, fecundity) and the environmental temperature (Fig. 1a) (Huey & Stevenson 1979; Angilletta 2009). Specifically, Huey & Kingsolver (1993) proposed four possible responses of TPC to selection on heat tolerance: *i*) in absence of genetic correlations between performance at high and low temperatures, the increase of CT_max_ might increase the TPC breadth (Fig. 1a); *ii*) if the thermal limits are negatively correlated, the selection for a higher CT_max_ might reduce the performance at low temperatures (e.g. lower thermal limit, CT_min_), then moving the entire TPC to higher temperatures (Fig. 1b); *iii*) if CT_max_ is positively correlated to the maximum performance, then the selection to high heat tolerance should simultaneously increase the optimal temperature and the maximum performance (“hotter is better”, Fig. 1c); and finally *iv*) if maximum performance is negatively correlated to TPC breadth, then selection for a higher CT_max_ could boost the performance at low temperatures (“jack-of-all-temperatures is a master of none”, Fig. 1d). Previous evidence shows that acute thermal assays provide a precise estimation of the heat tolerance and its genetic component (Rezende et al. 2011; Santos et al. 2012; Castañeda et al. 2019). Thus, it is reasonable to expect that selection for heat tolerance under different thermal stress intensities will result into different correlated responses of TPC.

**Figure 1.**
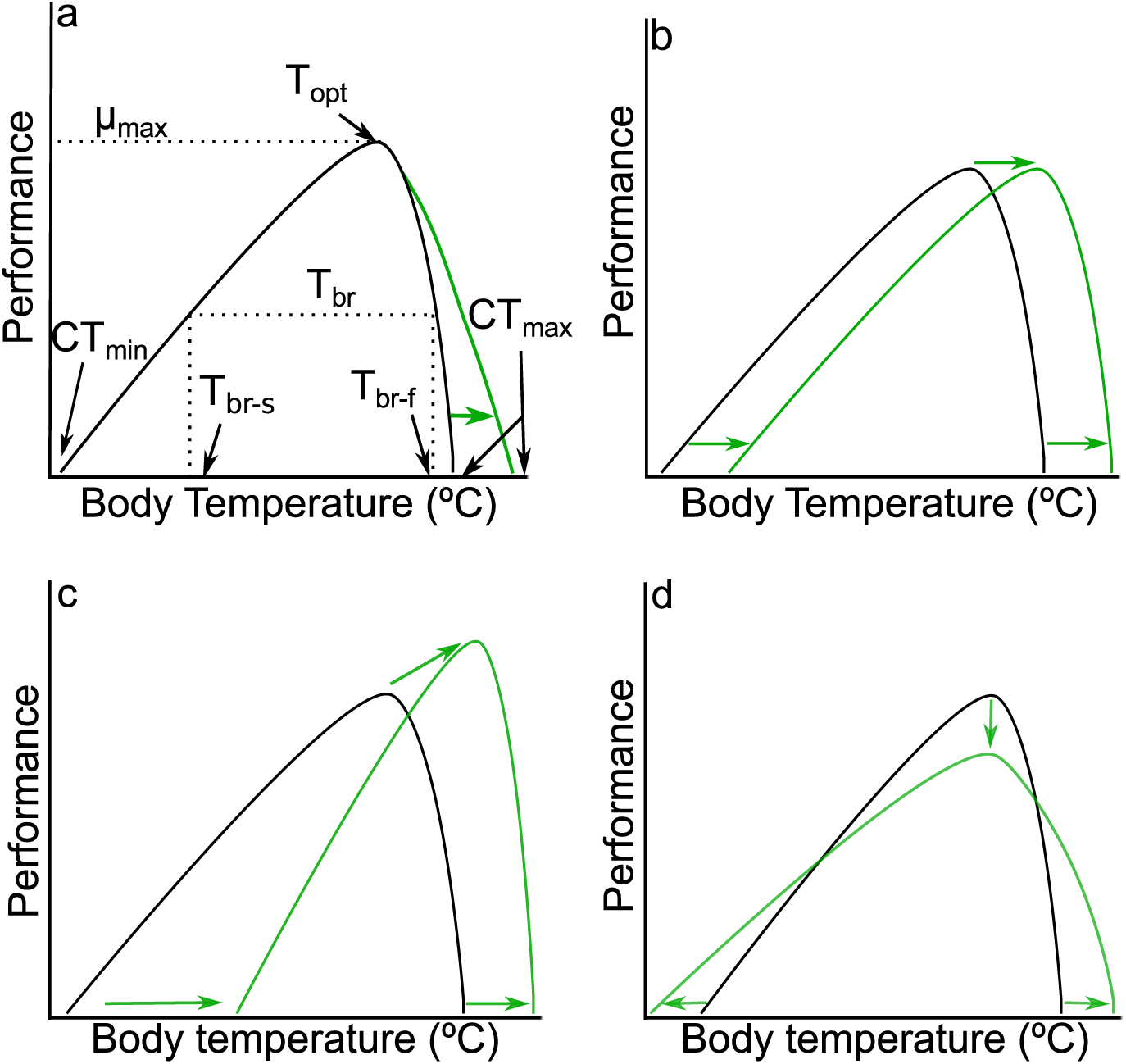
Thermal performance curve (TPC) and possible correlated responses to selection for increased heat tolerance (modified from Huey & Kingsolver 1993). a) Parameters of TPC and a hypothetical response of TPC when increased heat tolerance does not induce changes in other TPC parameters, b) increased heat tolerance leads to a displacement of TPC, c) increased heat tolerance increases µ_max_ and T_opt_ (hotter is better), and d) increased heat tolerance reduces µ_max_, but increases T_br_ (jack-of-all-temperatures is a master-of-none).

In the present work, we studied the impact of the thermal stress intensity on the evolutionary response of heat tolerance and the correlated response of TPC in *D. subobscura* (Collin). To accomplish this: (1) we compared the evolutionary response of heat tolerance to the artificial selection under two thermal intensity treatments: chronic (slow ramping selection, 0.08 ºC min^−1^) and acute thermal stress (fast ramping selection, 0.4 ºC min^−1^); and (2) we compared the correlated response of TPC to heat tolerance selection between different thermal intensity treatments (selected versus control lines), where TPC was estimated as the relationship between locomotor performance (i.e. climbing velocity) and environmental temperature. Because acute thermal assays provide a more precise estimation of heat tolerance and its genetic component (Rezende et al. 2011; Santos et al. 2012; Castañeda et al. 2019), we hypothesize that (1) the evolutionary response and realized heritability of the heat tolerance should be higher in selected lines under acute thermal stress than chronic thermal stress, and (2) as correlated responses should depend on the intensity of the thermal stress, we expect a correlated response of TPC only for selected lines under acute thermal stress compared to control lines and selected lines under chronic thermal stress. We used *D. subobscura* as a model organism because this species shows evidence of thermal adaptation in several phenotypic traits. For instance, latitudinal variation has been reported for body size, chromosomal inversion polymorphisms, desiccation resistance, thermal preference, heat tolerance and fertility (Budnik et al. 1991; Huey et al. 2000; Castañeda et al. 2013, 2015; Gilchrist et al. 2008; Porcelli et al. 2017). In addition, laboratory selection studies have demonstrated that populations of *D. subobscura* exposed for three years to different rearing temperatures show evolutionary responses in several traits such as chromosomal inversion polymorphisms, transcriptomic profiles, wing size, wing shape, and development and life-history traits (Santos et al. 2004, 2005, 2006; Laayouni et al. 2007;). Therefore, *D. subobscura* is a suitable model to evaluate the evolutionary responses of heat tolerance and TPC to high temperature selection.

## Methodology

### Fly maintenance

A mass-bred population of *D. subobscura* was established from the offspring of 100 isofemale lines derived from inseminated females collected in Valdivia, Chile (39.8 ºS 73.2 ºW). We dumped 10 females and 10 males from each isofemale line into an acrylic cage (27 × 21 × 16 cm^3^) to setup one large outbred population (>1500 breeding adults), which was maintained at 21 ºC (12:12 light:dark cycle) and feed with David’s killed-yeast *Drosophila* medium in Petri dishes (David 1962). At the next generation, a total of 150 eggs collected from Petri dishes were placed into 150-mL bottles with food, and a total of 45 bottles were maintained at 21 ºC. Emerged flies from 15 bottles were dumped into one acrylic cage, resulting in a total of three population cages: R1, R2 and R3. After three generations, with a larger population size and the environmental effects removed, each replicated cage (R1, R2 and R3) was split into four acrylic cages with more 1500 individuals per cage. The resulting four cages for each replicated cage were assigned to different treatments of our artificial selection experiment: acute-selected and acute-control lines, and chronic-selected and chronic-control lines. Thus, we established a total of 12 experimental lines, designated to four selection treatments where each treatment was replicated three times. All population cages were maintained on a nonoverlapping generation cycle and with controlled larval density as described in Castañeda et al. (2013, 2015).

### Selection on heat knockdown temperature

After three generations of the founding of experimental lines, three Petri dishes with David’s medium and extra dry yeast were placed into each experimental cage to collect eggs, which were transferred to vials with a density of 40 eggs per vial. Then, a total of 160 virgin females per experimental line were randomly chosen, individually placed into vials with fly medium, and mated with two non-related males from the same experimental line. After two days, males were discarded, and vials were checked for positive oviposition. This procedure was done before of the thermal selection assays because heat stress might cause mortality and sterilization in *Drosophila* (David et al., 2005). For each experimental line, 120 females were randomly chosen to evaluate the heat knockdown temperature as a proxy of CT_max_. Each female fly was individually placed in a capped 5-mL glass vial, which was attached to a rack with capacity to attach 60-capped vials (4 rows × 15 columns). Each rack was immersed in a water tank with an initial temperature of 28 ºC, which was controlled by a heating unit (Model ED, Julabo Labortechnik, Seelbach, Germany). After an equilibration time of 10 min, temperature was increased with a heating rate of 0.08 ºC min^−1^ for the chronic-selected lines or 0.4 ºC min^−1^ for the acute-selected lines. Each assay was photographed every 3 s with a high-resolution camera (D5100, Nikon, Tokyo, Japan). All photos for each assay were collated into a single file that was visualized to score the knockdown temperature defined as the temperature at which each fly ceased to move.

Knockdown temperatures were ranked, and we selected vials containing offspring of the 40 most tolerant female flies (upper 33% of each assay) to establish the next generation. From each one of these vials, four virgin females were individually placed in new vials with David medium and mated with two unrelated males. This procedure reestablished the original number of 160 female flies (4 females per vial × 40 vials = 160 females) before each heat thermal assay.

We performed the same procedure described above for control lines, except that founding flies were randomly chosen in each line. Briefly, the knockdown temperature of 40 females flies for each control line were assayed using a heating rate of 0.08 ºC min^−1^ for the chronic-control lines or 0.4 ºC min^−1^ for the acute-control lines. After measuring knockdown temperature for each female fly, we randomly selected 10 of them and we used their offspring to found the next generation of control lines.

Artificial selection for heat tolerance was performed for 16 generations, after which flies from each experimental line were dumped into an acrylic cage and maintained without selection at 21ºC (12:12 light-dark cycle), until the measurements of locomotor performance were performed at the generation 25.

### Locomotor performance

Locomotor performance was evaluated as the climbing velocity (cm s^−1^) in four days-old virgin females from generation 25. The locomotor performance for selected and control lines was measured at 5, 15, 20, 25, 30 and 35 ºC. For the experimental temperatures below to room temperature (~ 21 ºC), bath water was cooled using a cooling unit (Model FP 50-HP, Julabo Labortechnik, Seelbach, Germany), whereas for experimental temperatures above room temperature, bath water was heated using a heating unit (Model ED, Julabo Labortechnik, Seelbach, Germany). At each temperature, we evaluated 15 females from each experimental line, which were individually placed in 5 mL glass pipettes. The sealed pipettes were attached to a rack with a capacity of 15 pipettes, and the rack was immersed in a 40 L water tank setup to a specific experimental temperature. After 30 min in dark conditions, we lifted the rack 5 cm and let it fall to allow the flies dropped to the bottom of the pipettes, and then they climbed inside the pipettes. In addition, we turned on a light bulb placed over the water tank to induce the climbing taking advantage of positive phototropism and negative geotropism of flies. Each assay was video-recorded with a high-resolution camera (D5100, Nikon, Tokyo, Japan), and the video files were visualized to score the climbing velocity (cm s^−1^) for each assayed female.

## Statistical analysis

### Evolutionary response of upper thermal limit

We evaluated the evolutionary response of the knockdown temperatures by comparing the knockdown temperatures between selected and control lines at generation 16. We used a mixed linear model including the thermal selection as fixed factor and the replicated lines nested within thermal selection as random factor. This analysis was performed using the *lme* function of the *nlme* package for R (Pinheiro et al., 2018). As expected, we found that knockdown temperature was significant higher in acute-selected lines compared to chronic-selected lines (F_1,4_ = 1952.5, *P* = 1.6 × 10^−9^). Similar result was found for the comparison between control lines (F_1,4_ = 242.5, *P* = 8.1 × 10^−7^). Then, analyses were performed separately for the acute-selected and chronic-selected lines since it is known that acute stress assays estimate higher thermal tolerance than chronic stress assays (Chown et al. 2009; Castañeda et al. 2015, 2019).

We estimated the rate of evolutionary change for knockdown temperature in selected lines by calculating the Haldane index (*h*) for synchronic comparisons between selected and their respective control lines (see details in Hendry & Kinnison, 1999):

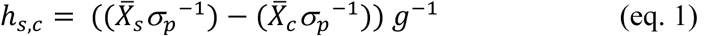

where 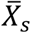 represents the mean knockdown temperature for each selected line and 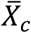 represents the pooled mean knockdown temperature between control lines, σ_*_ is the pooled standard deviation of selected and control lines, and *g* is the number of generations of thermal selection (*g* = 16 for the present study). For each thermal selection regime, we calculated the Haldane values for each replicated line (n = 3), and we evaluated if these values were significantly different from zero using a one sample t-test. Additionally, we compared whether the evolutionary rate of heat tolerance was different between both thermal selection treatments using a Kruskal-Wallis test.

The realized heritability (*h*^2^_n_) of the knockdown temperatures was calculated by regressing the cumulative response to selection against the cumulative selection differential. Because thermal selection was only performed on females and sires were randomly chosen, it is expected that the selection differential is only half. First, we estimated the cumulative response to selection on the cumulative selection differential for each replicated line, and these values were averaged within each selection treatment. Then, we regressed the averaged cumulative response to selection against the averaged cumulative selection differential forcing the regression through the origin, and the estimated slope was equated to the *h*^2^_n_ for each selection treatment (Walsh & Lynch, 2018). Finally, we compared the *h*^2^_n_ between selection treatments using a slope comparison approach: we used a linear model with the averaged cumulative response to selection as response variable and the interaction between the averaged cumulative selection differential and selection treatment as predictor variables. Differences of *h*^2^_n_, between selection treatments were considered when the interaction was significant.

### Locomotor performance

We measured the climbing velocity for the acute-and chronic-selected lines but because logistic reasons, we employed as control lines those lines belonging to the acute thermal stress control lines (Fig. S1). We estimated the TPC parameters for each replicated line (n = 9) by fitting four different models: the Lactin model (Lactin et al. 1995); the Briere model (Briere et al. 1999); the Performance model (Shi et al., 2011; Wang et al., 2013); and a modified Gaussian model. We found that the Lactin model showed the lowest Akaike Information Criteria (AIC) compared to the other three model (Table S1), so we use this model to estimate the TPC parameters: the optimal performance temperature (T_opt_), the maximum performance (μ_max_), the thermal breadth for the 50% and 80% of upper performance (T_br-50_ and T_br-80_, respectively), and the starting and ending temperatures of these thermal breadths (T_br-50s_, T_br-50f_, T_br-80s_ and T_br-80f_). Prior to TPC comparisons, normality and homoscedasticity of data were evaluated for all TPC parameters using the Lilliefors and Levene test, respectively (Table S2). Differences among TPC parameters were evaluated between acute-selected, chronic-selected and control lines using an one-way ANOVA. A posteriori differences among groups were evaluated using a HSD Tukey’s test. Only the T_br-80_ data were analyzed using the Kruskal-Wallis tests due to normality assumption violation. The TPC fitting was performed using the ThermPerf package for R (https://github.com/mdjbru-Rpackages/thermPerf).

## Results

### Evolutionary response of the upper thermal limit

Artificial selection for increasing heat tolerance resulted that the knockdown temperature evolved in both thermal selection treatments. We found that knockdown temperature was significant higher for the acute-selected lines in comparison to control lines (*X̄_acute_* ± SD = 37.71 ± 0.68°C and *X̄_control_* ± SD = 37.19 ± 0.90°C; F_1,4_ = 36.2, *P* = 0.004; Fig 2). Similarly, chronic-selected lines showed a significant increase of knockdown temperature in comparison to control lines (*X̄_chronic_* ± SD = 35.48 ± 0.66°C and *X̄_control_* ± SD = 34.97 ± 0.72°C; F_1,4_ = 41.7, *P* = 0.003; Fig. 2). On the other hand, we found no significant differences on the knockdown temperatures among replicated lines for acute thermal stress (χ^2^_2_ = 1.9 × 10^−7^, *P* = 0.99) neither for chronic thermal stress (χ^2^_2_ = 3.3 × 10^−7^, *P* = 0.99).

**Figure 2.**
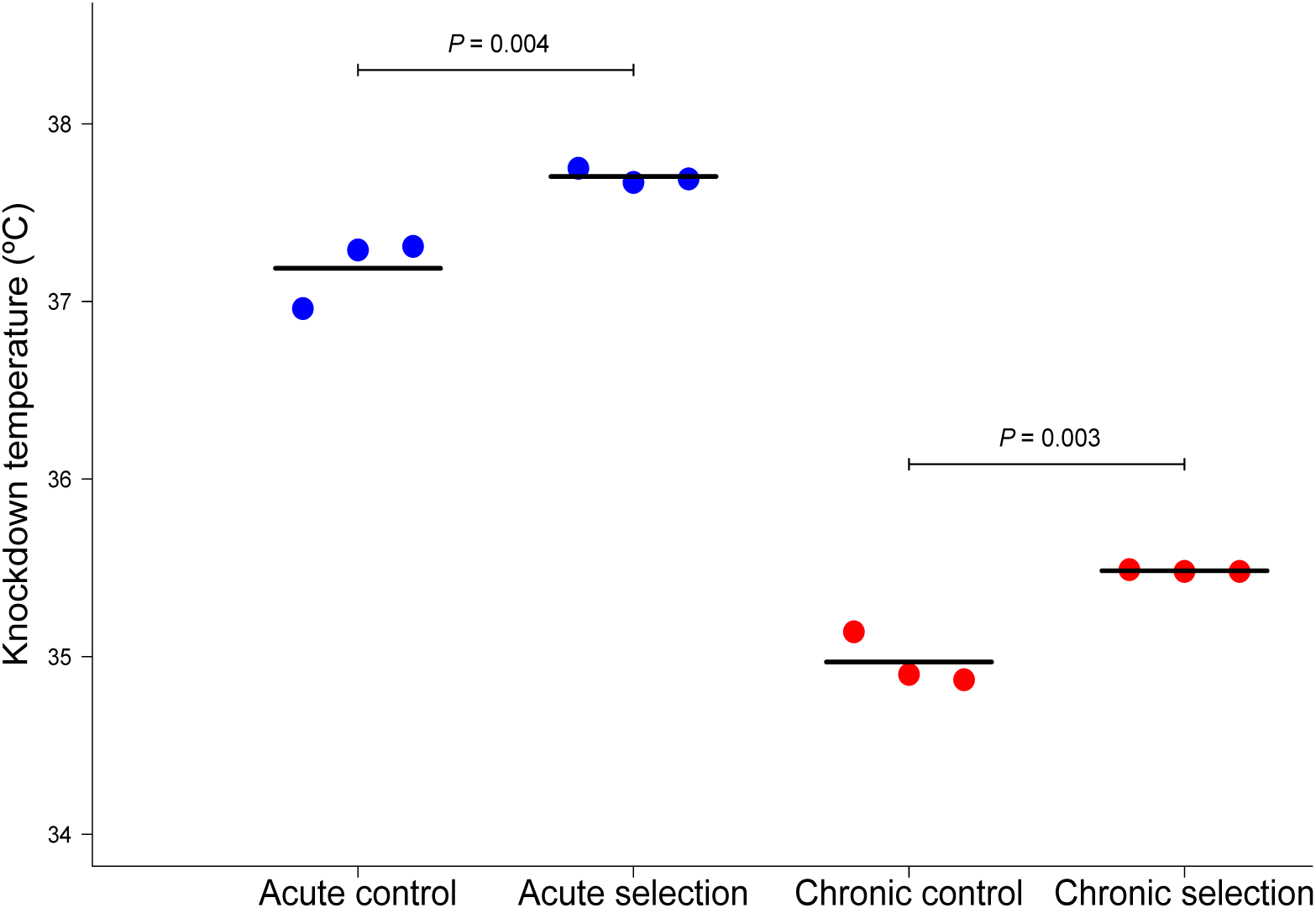
Knockdown temperature (ºC) of acute-control and acute-selected lines (blue) and chronic-control and chronic-selected lines (red) after 16 generations of artificial selection for increasing heat tolerance in *Drosophila subobscura*. Dots represent the mean value for each replicated line, and black lines represent the overall mean of each selection or control treatment. *P*-values were estimated using a mixed-linear model with selection treatment as fixed effect and replicated line as random effects nested within selection treatment.

We found that the heat tolerance showed a significant different evolutionary rate from zero, both for acute-selected (*h* acute,control ± SD = 0.061 ± 0.008, t_2_ = 13.30, *P* = 0.006) and chronic-selected lines (*h* chronic-control ± SD = 0.060 ± 0.004, t_2_ = 26.99, *P* = 0.001).

However, evolutionary rates were similar between both thermal selection treatments (Kruskal-Wallis, χ^2^_2_ = 2.0, *P* = 0.37). Additionally, the realized heritabilities of the knockdown temperatures were significantly different from zero for acute-selected (*h*^2^_pooled-replicates_ = 0.180 ± 0.013, t_1_ = 13.75, *P* < 0.0001) and chronic-selected lines (*h*^2^_pooled-replicates_ = 0.226 ± 0.020, t_1_ = 11.46, *P* < 0.0001). Despite the realized heritability was higher for chronic-selected lines, it was not significantly different from those estimated for acute-selected lines (F_1,20_ = 0.28, *P* = 0.60).

### TPC correlated evolution

Because locomotor performance was evaluated in at generation 25 (after nine generations without thermal selection on heat tolerance), we evaluated the knockdown temperature in all experimental lines before the climbing assays with the same protocol previously applied. Similarly to our findings found at generation 16, we found that knockdown temperatures of the acute-selected lines were significantly higher than those exhibited for the control lines (*X̄_acute_* ± S.D. = 38.59 ± 0.98°C and *X̄_control_* ± S.D. = 37.61 ± 1.45°C; F_1,4_ = 56.4, *P* = 0.002), and chronic-selected lines also showed a higher knockdown temperature than control lines (*X̄_chronic_* ± S.D. = 35.92 ± 0.67°C and *X̄_control_* ± S.D. = 35.14 ± 0.81°C; F_1,4_ = 83.5, *P* = 0.001).

We found that some TPC parameters were significantly influenced by the thermal selection regime (Table 1 for parameter estimated using the Lactin model. Table S3 and S4 show the same parameter estimated using the Performance and modified Gaussian models, respectively). For instance, T_opt_ was significantly influenced by thermal selection (*F*_2,6_ = 6.9, *P* = 0.02). Interestingly, acute-selected lines showed a higher T_opt_ than control lines (Table 1; Fig. 3; HSD Tukey: *P* = 0.02), chronic-selected lines exhibited a similar T_opt_ than control lines (Table 1; Fig. 3; HSD Tukey: *P* = 0.269), and acute-selected and chronic-selected lines did not differ between them (Table 1; Fig. 3; HSD Tukey: *P* = 0.197). Additionally, temperatures delimiting the thermal breadth for the 50% and 80% of upper performance (T_br-50_ and T_br-80_, respectively) were significantly affected by thermal selection Table 1; Fig. 3). Specifically, initial and final temperatures of T_br-50_ (Table 1) were significantly different between experimental lines (T_br-50s_: F_2,6_ =9.89, *P*=0.01; and T_br-50f_: F_2,6_ = 10.1, *P* = 0.01, respectively), similarly to initial and final temperature of T_br-80_ (T_br-80s_: F_2,6_ = 10.01, *P* = 0.01; and T_br-80f_: F_2,6_ = 10.02, *P* = 0.01). We found that delimiting temperatures were significantly higher for acute-selected lines than for control lines (HSD Tukey: T_br-50s_: *P* = 0.01; T_br-50f_: *P* = 0.01; T _br-80s_: *P* = 0.01; T _br-80f_: *P* = 0.01). Whereas, these parameters were similar between chronic-selected and control lines (HSD Tukey: T_br-50s_: *P* = 0.231; T_br-50f_: *P* = 0.23; T _br-80s_: *P* = 0.204; T_br-80f_: *P* =0.205), and they also were similar between both thermal selected lines (HSD Tukey: T_br-50s_: *P* = 0.093; T_br-50f_: *P* = 0.088; T_br-80s_: *P* = 0.101; T_br-80f_: *P*= 0.1). Finally, the maximum performance (μ_max_) was not significantly affected by the thermal selection (F_2,6_ = 1.77, *P* = 0.25), similarly to the response exhibited by the thermal breadth for the 50% (F_2,6_ = 0.64, *P* = 0.56) and 80% of upper performance (F_2,6_ = 0.80, *P* = 0.49). Summarizing, our artificial selection experiment for increasing heat tolerance of *D. subobscura* resulted in a correlated response on the TPC, but only for the acute-selected lines. Therefore, an evolutionary increase of the heat tolerance induced a displacement of TPC to higher temperatures without changes on its topology (Fig. 3).

**Table 1.** Parameters (mean ± S.E.) of the thermal performance curve (estimated as climbing velocity) estimated with the Lactin model in acute-selected, chronic-selected and control lines in an artificial selection experiment for increasing heat tolerance in *Drosophila subobscura*.

|  | Acute-selected lines | Chronic-selected lines | Control lines |
| --- | --- | --- | --- |
| $T_{opt}$ ($^{\circ}\text{C}$) | 30.08 ± 0.73 | 29.22 ± 0.52 | 28.46 ± 0.24 |
| $\mu_{max}$ ($\text{cm s}^{-1}$) | 0.80 ± 0.08 | 0.68 ± 0.07 | 0.80 ± 0.12 |
| $T_{br-50}$ ($^{\circ}\text{C}$) | 14.99 ± 0.01 | 14.99 ± 0.01 | 14.98 ± 0.02 |
| $T_{br-50s}$ ($^{\circ}\text{C}$) | 19.12 ± 0.15 | 18.78 ± 0.31 | 18.32 ± 0.16 |
| $T_{br-50f}$ ($^{\circ}\text{C}$) | 34.11 ± 0.15 | 33.77 ± 0.31 | 33.30 ± 0.16 |
| $T_{br-80}$ ($^{\circ}\text{C}$) | 5.99 ± 0.01 | 5.99 ± 0.01 | 5.99 ± 0.01 |
| $T_{br-80s}$ ($^{\circ}\text{C}$) | 26.25 ± 0.26 | 25.71 ± 0.48 | 25.01 ± 0.22 |
| $T_{br-80f}$ ($^{\circ}\text{C}$) | 32.24 ± 0.26 | 31.70 ± 0.48 | 30.99 ± 0.22 |

**Figure 3.**
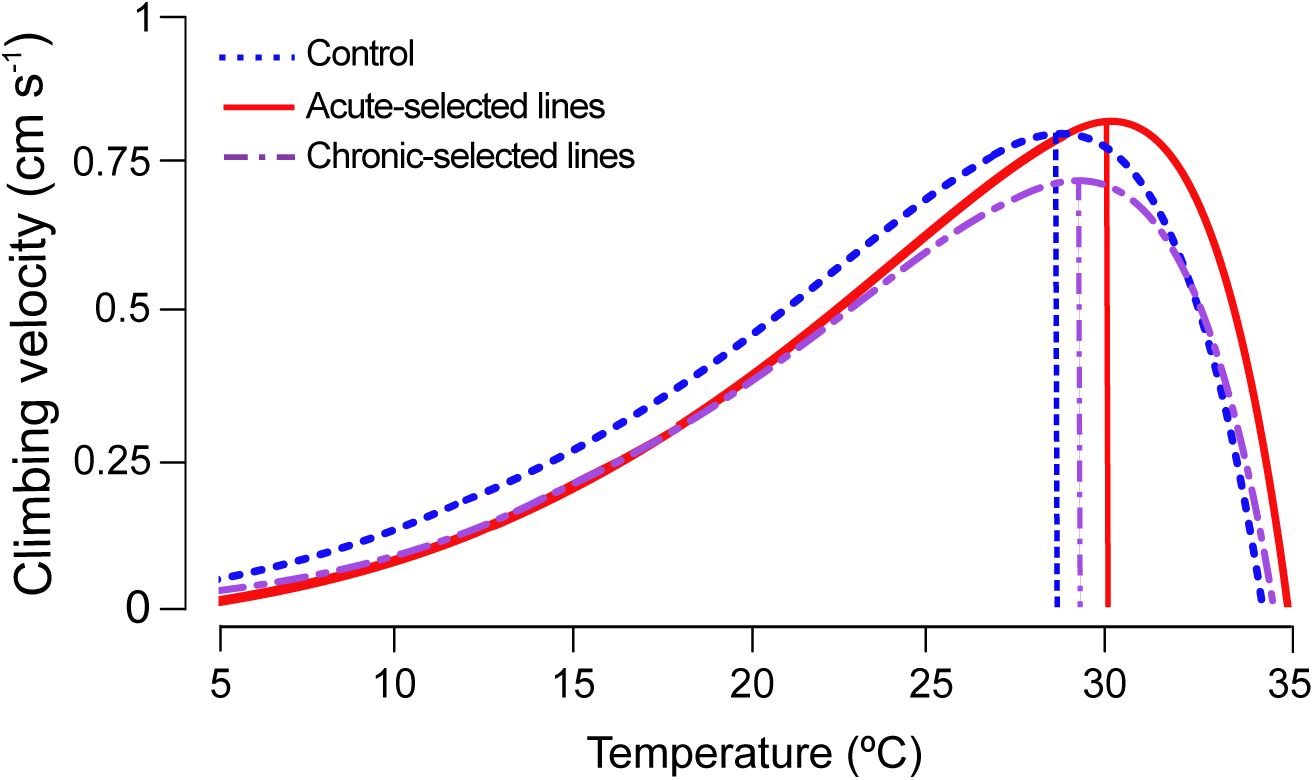
Thermal performance curves estimated from the climbing velocity measured at 6 temperatures (5, 15, 20, 25, 30 and 35 ºC) for acute-selected (red), chronic-selected (purple) and control lines (blue) from an artificial selection experiment for increasing heat tolerance in *Drosophila subobscura*. Vertical lines represent the averaged optimal temperature (n = 3) for each selection treatment. Climbing velocity for each replicated line of each selection treatments can be visualized in the Fig. S1.

## Discussion

When evaluating the effects of global warming on distributions and survival of ectotherm organisms becomes essential to contrast and predict the evolutionary responses of the thermal limits across populations (Panetta et al., 2018). In the present study, we evaluated the evolutionary response of the upper thermal limit in *D. subobscura* populations selected for an increasing heat tolerance at different thermal stress intensities. We demonstrated that upper thermal limits of *D. subobscura* evolved independently of thermal stress intensity, suggesting that thermal tolerance to high temperatures has enough genetic variation to respond to an increase in temperatures due to global warming. Our results are particularly interesting because the arrival of *D. subobscura* to South America from Europe was characterized by a strong founder effect due to reduced number of colonizing individuals, resulting in a reduced genetic variation before the expansion along the colonized geographical range in Chile (Pascual et al., 2007). Despite that, *D. subobscura* exhibits signals of thermal adaption only a few decades after of its introduction in South America (Brncic et al. 1981; Ayala et al., 1989; Huey et al., 2000; Castañeda et al., 2013, 2015). Additionally, the evolutionary response of heat tolerance resulted after artificial thermal selection performed only in females, which should reduce the selection effectiveness near to 50% (Huey & Kingsolver, 1993).

The evolution of the upper thermal limit reported in this work agrees with previous increases of CT_max_ in experiments of artificial selection in *D. melanogaster* and *D. buzzatii* (Bubliy and Loeschcke, 2005; Folk et al., 2006; Sambucetti et al., 2010; Hangartner and Hoffmann et al., 2016), but contrasts with the multiple evidences of a limited evolutionary potential to heat tolerance in the *Drosophila* clade (Chown et al., 2009; Mitchell and Hoffmann, 2012; Kellerman et al., 2012; Blackburn et al., 2014; van Heerwaarden et al., 2015; MacLean et al., 2019). In this sense, unlike studies of artificial selection, comparatives studies provide with a wide vision of thermal evolution in the *Drosophila*, suggesting that the evolution of heat tolerance is severely constrained (Kellerman et al., 2012; MacLean et al., 2019). However, it has been widely demonstrated that the capacity for detecting evolutionary patterns of CT_max_ in intra-and interspecific studies can be hindered by uncontrolled factors as the intensity of thermal stress because chronic thermal stress decreases the accuracy to detect these patterns (Rezende et al., 2014; Chown et al., 2009; Mitchell and Hoffmann 2010; Blackburn et al., 2014; Kingsolver and Umbanhowar, 2018; Sunday et al., 2019; Castañeda et al., 2019; Kovacevic et al., 2019). In fact, genetic quantitative studies in *Drosophila* have demonstrated that long thermal assays (e.g. chronic stress or slow ramping assays) provide a lower estimations of the evolutionary potential for heat tolerance compared to short thermal assays (e.g. acute stress or fast ramping assays). This is probably because long assays increase the environmental variance of heat tolerance and then, heritability estimates become lower (Chown et al., 2009; Mitchell and Hoffmann 2010; Blackburn et al., 2014; Castañeda et al. 2019). According to this, the evolutionary capacity of heat tolerance should depends on the intensity of thermal stress thus we expected a reduced response of the heat tolerance when the artificial selection was performed using a chronic thermal stress. Despite these predictions, we did not find significant differences in the evolutionary change rate or realized heritability of heat tolerance between thermal selection treatments (averaged *h*^2^n = 0.18 for acute-selected lines, and averaged *h*^2^n = 0.23 for chronic-selected lines). These realized heritabilities are closer to the mean heritability reported for terrestrial ectotherms (*h*^2^ = 0.28; Diamond, 2017), whereas the heritability estimated for chronic-selected lines was even higher than those previously reported in studies of thermal selection or using quantitative genetics (Gilchrist et al., 1999; Mitchell & Hoffmann, 2010; Blackburn et al., 2014). In a previous work, our group estimated the narrow-sense heritability (*h*^2^) for heat tolerance in *D. subobscura* using static (38 ºC) and ramping assays (0.1 ºC min^−1^) (Castañeda et al., 2019). We found than *h*^2^ using static assay was higher (*h*^2^ = 0.134) that those estimated using a slow ramping assay (*h*^2^ = 0.084), supporting the hypothesis that acute thermal assays provide a more precise estimation of the genetic component of heat tolerance in comparison to chronic thermal assays.

We performed artificial selection on heat tolerance for 16 generations and for logistic reasons, we were able to perform the climbing assays nine generations later. This means that selected lines experience relaxed thermal selection during nine generations. Hence, we checked for differences of heat tolerance between selected and control lines before performing the climbing assays. A plausible result of this analysis could have been that relaxed selection allows a reversion of the evolutionary increase of heat tolerance in selected populations because the high energy costs associated with an increase of thermotolerance (Callahan et al., 2008; Lahti et al., 2009). Previous evidence indicates that selection on heat tolerance results into an increase in heat-shock protein (HSP) levels, which is a well-known thermoprotective mechanism that maintain the cellular homeostasis (Sørensen et al., 1999). However, the increase of the HSP production leads to an increase of the metabolic demand (Hoekstra & Montooth, 2013) and reduction of the reproductive output (Krebs & Loeschcke, 1994; Krebs & Feder, 1997). Thus, if thermal selection would have led to elevated energy costs associated with an increase of thermotolerance, a reversion of the upper thermal limit should be expected after the relaxation of thermal selection. However, we found that the evolutionary response of the upper thermal limit was maintained after nine generations without thermal selection, suggesting that the evolutionary increase of the heat tolerance in *D. subobscura* did not involve an increase of maintenance costs. These results agree with previous studies in *D. melanogaster* (Williams et al., 2012) and in the copepod *Trigriopus californicus* (Kelly et al., 2013), in which the evolution of heat tolerance was principally limited by the presence of genetic variation, showing no tradeoff or cost that limit their evolutionary responses. Moreover, we also found that metabolic rate did not differ between selected and control lines and moreover, fecundity increased in the selected lines compared to control lines (A. Mesas, unpublished results).

Evolutionary theory predicts that thermal performance of ectotherms should evolve in response to environmental temperatures (Huey and Kingsolver, 1989; Angilletta, 2009). High environmental temperatures are expected to impose selective pressures on natural populations, resulting in evolution of heat tolerance and TPC (Huey & Kingsolver, 1993). To our knowledge, the present work shows for first time the evolution of TPC as response to artificial selection on CT_max_, suggesting that performance at low and high temperatures could be negatively correlated. Additionally, the evolutionary response of TPC depended on thermal stress intensity employed during thermal selection. Specifically, we found a displacement of the TPC to higher temperatures only in the acute-selected lines, suggesting that associations between different TPC parameters are more likely to be detected using acute thermal assays, especially for heritability estimates (Mitchell and Hoffmann, 2010; Blackburn et al., 2014; van Heerwaarden & Sgrò, 2014; Castañeda et al., 2019). This TPC displacement to high temperatures is according with one of the scenarios proposed by Huey and Kingsolver (1993), in contrast to other two well-supported scenarios such as “hotter is better” (Angilletta et al., 2010; Nati et al., 2016) and “jack-of-all-temperatures is a master-of-none” (Huey & Hertz 1984; Gilchrist, 1995). Our results also agree with previous results of thermal selection experiments in *Drosophila* species (Huey & Kingsolver, 1993; Bubliy & Loeschcke, 2005; Mori & Kimura, 2008; Diamond et al., 2017) and in ectotherm vertebrates as *Anolis sagrei* (Logan et al., 2014), suggesting that the natural selection mediated by high temperatures is strong enough to produce changes in thermal performance of ectotherm species (but see Gilbert & Miles, 2017).

In conclusion, the intensity of thermal stress has important effects on the evolution of thermal physiology in ectotherms, indicating that different thermal scenarios might yield a similar evolution for heat tolerance but having contrasting consequence on thermal performance. Therefore, the intensity of thermal stress could also have important consequences on the estimations of vulnerability of ectotherms to global warming.

## Acknowledgments

We thank Marcela Morales for her work to establish the experimental lines and Sergio Estay for his methodological advices to compare thermal performance curves. Roberto Nespolo and Mauro Santos provided helpful comments on this manuscript. AM thanks a CONICYT fellowship no. 21140595 (Chile) and a Postdoctoral fellowship from Universidad del Bío-Bío, Decreto Exento RA N° 352/11510/2019 (Chile). This work was funded by the FONDECYT grant 1140066 (Chile) to LEC.

## Author contribution

AM designed experiments and conducted experiments and statistical analyses, and wrote the manuscript. AJ maintained the experimental animals, performed experiments and approved the final version of the manuscript. LEC conceived the original idea, designed experiments, conducted statistical analyses, provided funds for all experiments, and heavily edited and wrote the manuscript.

