## Supplementary Material for "Correlated evolution between heat tolerance and thermal performance curves in *Drosophila subobscura*"

**Table S1**. TPC models fitted for selected and control lines ………………………..……..… 2

**Table S2**. Normality and homoscedasticity test for TPC paraments …….………….…..…… 4

**Table S3**. TPC paraments estimated using the Performance model ..…….……………..…… 5

**Table S4**. TPC paraments estimated using the modified Gaussian model …..…………..…… 6

**Fig. S1**. TPC models fitted for selected and control lines ……………..……………..…… 7

**Appendix S1**. Functions to describe TPC models ………………………………………… 8

**Table S1**. Comparison of models used to describe the thermal performance curves (measured as climbing velocity at 6 different temperatures) in *Drosophila subobscura*. Acute-selected, chronic-selected and control lines correspond to different artificial selection experiments for increasing heat tolerance in *D. subobscura*. Models (Breire, Gaussian, Lactin and Performance models) were fitted to each replicated lines (R1, R2 and R3) belong to the different selection treatments and were ordered from lowest to highest corrected Akaike Information Criteria (AICc) value. Lactin model (in bold) always showed the best fit (lowest AICc value) for all replicated lines and for this reason, TPC parameters were estimated using this model.

| | Replicates | Models | AICc | ΔAICc | Weight | $R^2$ |
|---|---|---|---|---|---|---|
| Acute-selected lines | R1 | **Lactin** | **21.196** | **0.000** | **9.3E-01** | **0.600** |
| | | Breire | 26.398 | 5.202 | 6.9E-02 | 0.567 |
| | | Performance | 32.777 | 11.581 | 4.04E-03 | 0.548 |
| | | Gaussian | 54.340 | 33.145 | 5.9E-08 | 0.409 |
| | R2 | **Lactin** | **1.685** | **0.000** | **8.3E-01** | **0.585** |
| | | Breire | 4.806 | 3.121 | 1.7E-01 | 0.561 |
| | | Performance | 6.353 | 4.668 | 1.02E-01 | 0.566 |
| | | Gaussian | 29.145 | 27.460 | 9.0E-07 | 0.425 |
| | R3 | **Lactin** | **37.281** | **0.000** | **5.2E-01** | **0.576** |
| | | Breire | 37.453 | 0.172 | 4.8E-01 | 0.567 |
| | | Performance | 44.189 | 6.908 | 2.3E-02 | 0.546 |
| | | Gaussian | 57.948 | 20.667 | 1.7E-05 | 0.456 |
| Chronic-selected lines | R1 | **Lactin** | **26.051** | **0.000** | **1.0E+00** | **0.509** |
| | | Performance | 31.737 | 5.686 | 7.6E-02 | 0.481 |
| | | Gaussian | 38.737 | 12.685 | 1.8E-03 | 0.422 |
| | | Breire | | Did not converged | | |

| | | | | | | |
|---|---|---|---|---|---|---|
| | R2 | **Lactin** | **14.901** | **0.000** | **9.9E-01** | **0.570** |
| | | Performance | 16.758 | 1.857 | 3.6E-01 | 0.565 |
| | | Gaussian | 24.751 | 9.851 | 7.2E-03 | 0.510 |
| | | Breire | | did not converged | | |
| | R3 | **Lactin** | **32.962** | **0.000** | **6.6E-01** | **0.376** |
| | | Performance | 33.802 | 0.84 | 3.8E-01 | 0.375 |
| | | Breire | 34.267 | 1.306 | 3.4E-01 | 0.353 |
| | | Gaussian | 45.665 | 12.703 | 1.1E-03 | 0.266 |
| Control lines | R1 | **Lactin** | **50.303** | **0.000** | **9.8E-01** | **0.568** |
| | | Performance | 56.170 | 5.867 | 6.9E-02 | 0.543 |
| | | Gaussian | 58.255 | 7.951 | 1.8E-02 | 0.518 |
| | | Breire | | Did not converged | | |
| | R2 | **Lactin** | **43.141** | **0.000** | **8.2E-01** | **0.564** |
| | | Performance | 45.819 | 2.678 | 2.3E-01 | 0.564 |
| | | Breire | 46.160 | 3.019 | 1.8E-01 | 0.559 |
| | | Gaussian | 55.091 | 11.950 | 2.1E-03 | 0.502 |
| | R3 | **Lactin** | **41.982** | **0.000** | **7.9E-01** | **0.438** |
| | | Performance | 43.159 | 1.177 | 3.8E-01 | 0.435 |
| | | Gaussian | 44.597 | 2.615 | 2.1E-01 | 0.408 |
| | | Breire | | Did not converged | | |

**Table S2**. Tests of normality (Lilliefors test) and homoscedasticity (Levene test) for thermal performance curve (TPC) parameters estimated with Lactin model to climbing velocity of *D. subobscura*. TPC parameters were estimated for acute-selected, chronic-selected and control lines from an artificial selection experiment for increasing heat tolerance in *Drosophila subobscura*.

| | Lilliefors test | | Levene test | | |
|---|---|---|---|---|---|
| | D value | P-value | DF | F Value | P-value |
| $T_{opt}$ | 0.19898 | 0.384 | 2,6 | 0.5363 | 0.6105 |
| $\mu_{max}$ | 0.23723 | 0.1525 | 2,6 | 0.0632 | 0.9394 |
| $T_{br\text{-}50}$ | 0.24506 | 0.1234 | 2,6 | 1.3889 | 0.3194 |
| $T_{br\text{-}50s}$ | 0.20844 | 0.3129 | 2,6 | 0.5841 | 0.5865 |
| $T_{br\text{-}50f}$ | 0.20216 | 0.3591 | 2,6 | 0.5993 | 0.579 |
| $T_{br\text{-}80}$ | 0.33332 | 0.0047 | 2,6 | 0.3333 | 0.729 |
| $T_{br\text{-}80s}$ | 0.21146 | 0.292 | 2,6 | 0.5936 | 0.5818 |
| $T_{br\text{-}80f}$ | 0.20961 | 0.3047 | 2,6 | 0.5885 | 0.5843 |

**Table S3**. Parameters (mean ± S.E.) of the thermal performance curve (measured as climbing velocity) estimated with the Performance model for acute-selected, chronic-selected and control lines from an artificial selection experiment for increasing heat tolerance in *Drosophila subobscura*.

| **Performance** | | | | | | | | | |
|---|---|---|---|---|---|---|---|---|---|
| | Acute-selected lines | | | Chronic-selected lines | | | Control lines | | |
| $T_{opt}$ (ºC) | 33.01 | ± | 1.22414 | 30.8433 | ± | 2.381645 | 28.75 | ± | 1.12743 |
| $\mu_{max}$ (cm $s^{-1}$) | 0.829 | ± | 0.03969 | 0.66276 | ± | 0.044052 | 0.77034 | ± | 0.12003 |
| $T_{br\text{-}50}$ (ºC) | 15 | ± | 0 | 15 | ± | 0 | 15 | ± | 0 |
| $T_{br\text{-}50s}$ (ºC) | 20.62 | ± | 0.36373 | 19.0133 | ± | 19.01333 | 18.25 | ± | 0.47634 |
| $T_{br\text{-}50f}$ (ºC) | 35.62 | ± | 0.36373 | 34.0133 | ± | 0.684129 | 33.25 | ± | 0.47634 |
| $T_{br\text{-}80}$ (ºC) | 6 | ± | 0 | 6 | ± | 0 | 6 | ± | 0 |
| $T_{br\text{-}80s}$ (ºC) | 29.06 | ± | 0.88047 | 26.7066 | ± | 1.567206 | 25.1666 | ± | 0.94769 |
| $T_{br\text{-}80f}$ (ºC) | 35.06 | ± | 0.88047 | 32.7066 | ± | 1.5672 | 31.1666 | ± | 0.94769 |

**Table S4**. Parameters (mean ± S.E.) of the thermal performance curve (measured as climbing velocity) estimated with the modified Gaussian model for acute-selected, chronic-selected and control lines from an artificial selection experiment for increasing heat tolerance in *Drosophila subobscura*.

| **Gaussian** | Acute-selected lines | | | Chronic-selected lines | | | Control lines | | |
|---|---|---|---|---|---|---|---|---|---|
| $T_{opt}$ (ºC) | 25.673 | ± | 0.14011 | 25.356 | ± | 0.18770 | 25.0566 | ± | 0.1184 |
| $\mu_{max}$ (cm $s^{-1}$) | 0.6297 | ± | 0.06171 | 0.5707 | ± | 0.10106 | 0.73142 | ± | 0.0990 |
| $T_{br-50}$ (ºC) | 15.003 | ± | 0.00577 | 15 | ± | 0 | 15 | ± | 0 |
| $T_{br-50s}$ (ºC) | 17.276 | ± | 0.14571 | 16.953 | ± | 0.19347 | 18.3933 | ± | 3.1165 |
| $T_{br-50f}$ (ºC) | 32.28 | ± | 0.15132 | 31.953 | ± | 0.19347 | 30.3933 | ± | 2.0839 |
| $T_{br-80}$ (ºC) | 6.0033 | ± | 0.00577 | 6 | ± | 0 | 5.99666 | ± | 0.0057 |
| $T_{br-80s}$ (ºC) | 22.533 | ± | 0.14011 | 22.206 | ± | 0.17925 | 18.3933 | ± | 2.9242 |
| $T_{br-80f}$ (ºC) | 28.556 | ± | 0.12423 | 28.206 | ± | 0.17925 | 30.3966 | ± | 2.2747 |

**Figure S1**. Climbing velocity measured at 6 temperatures (5, 15, 20, 25, 30 and 35 ºC) for flies belong to control (blue), acute-selected (red), and chronic-selected (purple) lines from an artificial selection experiment for increasing heat tolerance in *Drosophila subobscura*. Thermal performance curves were estimated using Briere, modified Gaussian, Lactine and Performance (different lines in each plot). According to the corrected Akaike information criteria (AICc), Lactin model (green lines) always showed the best fit (lowest AICc value, Table S1) for all replicated lines.

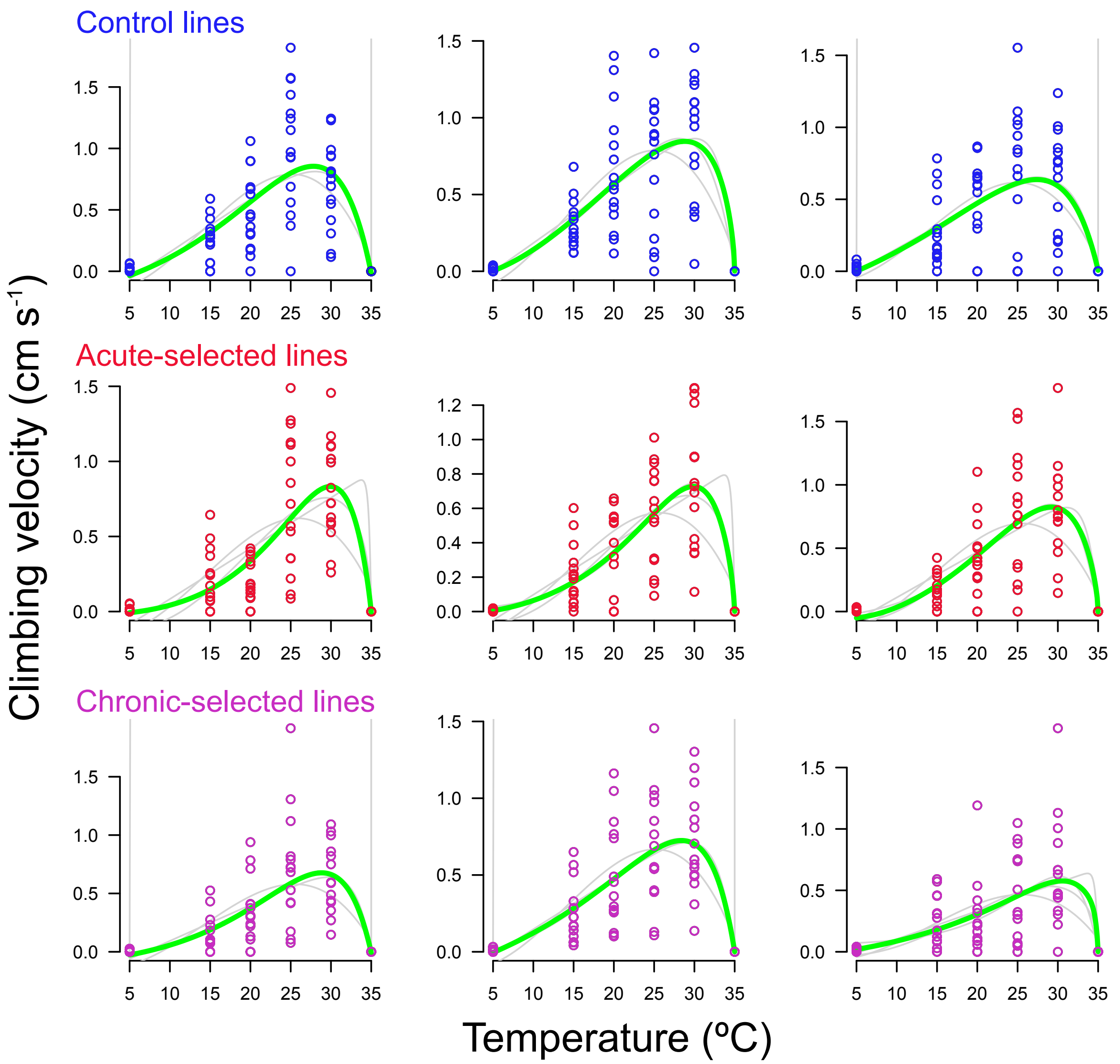

**Appendix S1**. Functions to describe the relationship between temperature (T$^{o}$) and performance

*Briere model*

Thermal Performance= $a\ T^{o}\ (T^{o}-CT_{min})(CT_{max}-T^{o})^{1/2}$

*Lactin model*

Thermal Performance= $b + \exp(CT_{min}\ T^{o}) – \exp(CT_{min}\ CT_{max} –(CT_{max}-T^{o})/d)$

*Modified Gaussian model*

Thermal Performance= $a + b\ T^{o2} \log (T^{o}) + c\ T^{o3}$

*Performance model*

Thermal performace= $c\ (T^{o}\ CT_{min})\ 1 – \exp\ (k(T^{o}-CT_{max}))$
